# Bidirectional communication between neurons in the mesentery and ileal myenteric neurons

**DOI:** 10.64898/2026.07.04.736073

**Authors:** Pieter Vanden Berghe, Feifei Guo, Klaas Van Mechelen, Zhiling Li, Candice Fung

**Affiliations:** Laboratory for Enteric NeuroScience (LENS), Translational Research in Gastrointestinal Disorders (TARGID), KU Leuven, Leuven, Belgium; Physiology & Pathophysiology Department, School of Basic Medicine, Qingdao University; Qingdao, China

**Keywords:** extrinsic nerves, small intestine, mesentery, enteric nervous system, calcium imaging

## Abstract

The intestinal mesentery has been recently classified as a ‘new’ organ and contains various cell types including adipocytes, preadipocytes, endothelial cells, and immune cells. In addition, neuronal cell bodies are found in the small intestinal mesentery and are situated either individually or clustered together with glial cells in small ganglion structures close to the gut wall. However, little is known about the origin or function of these extra-intestinal mesenteric neurons. The aim of this study was to better these characterize mesenteric neurons and to examine their connectivity with the ENS using calcium imaging in adult mouse ileum with the mesentery attached. Here we show that neurons in the mesentery express typical ENS neurochemical markers, respond to 5-HT, ATP and the nicotinic agonist DMPP, and receive nicotinic synaptic inputs. Furthermore, using labeling with the neuronal tracer DiI, some mesenteric neurons were found to project into the gut wall and can provide functional excitatory inputs to myenteric neurons. By contrast, we did not find evidence for mesenteric neurons providing inputs to other extrinsic neuronal targets, suggesting that they preferentially interact with the ENS. We also demonstrate that mesenteric neurons can be activated by intestinal distension and that the mesentery provides a source of inhibition to the myenteric plexus. Taken together, we show that the ENS not only interacts with vagal and spinal afferents, and sympathetic and parasympathetic nerves, but also neurons situated in the mesentery. Finally, our data suggest that these neurons may provide a form of negative feedback to the myenteric plexus such as in the event of intestinal distension. These findings have important implications for the regulation of intestinal motility in physiological and pathophysiological conditions.

## Introduction

The mesentery was long considered to be a collection of fragmented connective tissues linking digestive organs to the abdominal cavity. However, it has been recently found to be a continuous tissue in or on which the digestive organs form during embryonic development, and they remain anatomically connected through to adulthood. Thus, the mesentery has been re-classified as a ‘new’ organ (Coffey et al., 2022; Coffey & O’Leary, 2016) and emerging view is that it plays an active role in pathophysiological conditions (e.g. Crohn’s disease and inflammatory bowel diseases) (Lu et al., 2025; Yin et al., 2021). The mesentery comprises myriad cell types such as adipocytes, preadipocytes, endothelial cells, macrophages, immune cells, stromal cells (Schäffler et al., 2005), and notably, a small population of neurons, which are present as single neurons or are clustered in small ganglia situated close to the gut wall (Cracco & Filogamo, 1993; Yu et al., 2021). However, the function of these mesenteric neurons and their projection patterns are unknown. Their cell bodies are mainly found along axon bundles of mixed nerve types meandering through the mesentery, which project into or out of the gut wall. These include vagal and spinal afferent fibers, preganglionic parasympathetic and postganglionic sympathetic fibers that enter the intestine (Fung & Vanden Berghe, 2020), as well as fibers from enteric intestinofugal neurons, which project outside of the gut (Furness, 2003). These nerve bundles in turn run alongside blood and lymph vessels, and together they interconnect the gut and other organ systems. Whether mesenteric neurons interact with the enteric nervous system (ENS) and/or other extrinsic neuronal populations is unclear.

One of the earliest descriptions of mesenteric neurons was by Sheehan (1933), in which the author presented drawings of an unmyelinated nerve network with ganglion cells and fibers that coursed around the vasculature in cat and human mesentery. Since then, mesenteric neurons associated with rat ileum have also been reported (Cracco & Filogamo, 1993). More recently, Yu and colleagues (2021) described, in developing mouse embryos, a population of mesenteric neural crest cells (NCCs) distinct from vagal NCC. The authors proposed that these mesenteric NCCs enter and migrate into the gut wall to also contribute to ENS formation. Some of these mesenteric NCCs were shown to give rise to neurons and glia which are observed in the early postnatal (P7) gut.

In this study, we confirm that mesenteric neurons are present in small intestinal mesentery of mice and that they persist into adulthood. Using calcium imaging we show that these mesenteric neurons were functionally active, and that they send signals to and receive signals from myenteric neurons within the gut wall. These neurons appear to preferentially interact with the ENS rather than other extrinsic neuronal populations. Finally, we demonstrate that mesenteric neurons can respond to intestinal distension and that the mesentery provides a level of inhibition to the myenteric neuronal network. Collectively, these findings have important implications for the control of intestinal motility in physiology and pathophysiology.

## Results

### Neurons are found in the mesentery

Neurons were observed in the mesentery running alongside the full length of the small and large intestine. For this study, we focused on ileal region where we first identified these neurons and where they have been previously studied in rat (Cracco & Filogamo, 1993). We quantified the number of neurons and ganglia in the mesenteric bundle supplying the terminal ileum by immunolabelling with the pan-neuronal marker HuCD in tissue from Wnt1-Cre;R26R-GCaMP3 mice (Figure 1). In 6 week old mice, each mesenteric preparation contained a total of 219 ± 20 neurons (n = 4 preparations, N = 4 mice). These were present as individual neurons, in pairs, or clustered in ganglia of three or more neurons. Each preparation contained between 25 ± 2 ganglia, with each ganglion comprising 3-10 neurons. Mesenteric neurons and ganglia were mostly situated close to the gut wall, with 97 ± 1% of neurons found located around the vicinity of the tertiary branches of vasculature supplying the gut. By contrast, few neurons were found along the primary and secondary branches of vasculature (ranging from 0 – 7 neurons; 2 ± 1% of total neurons). We also examined the distribution of mesenteric neurons and ganglia along the length of the ileal loop (by dividing the mesentery into 30° sectors as shown in Figure 1A). The number of neurons and ganglia in the mesentery appear to increase from the proximal to the distal end of the ileum (Figure 1B-C).

**Figure 1.**
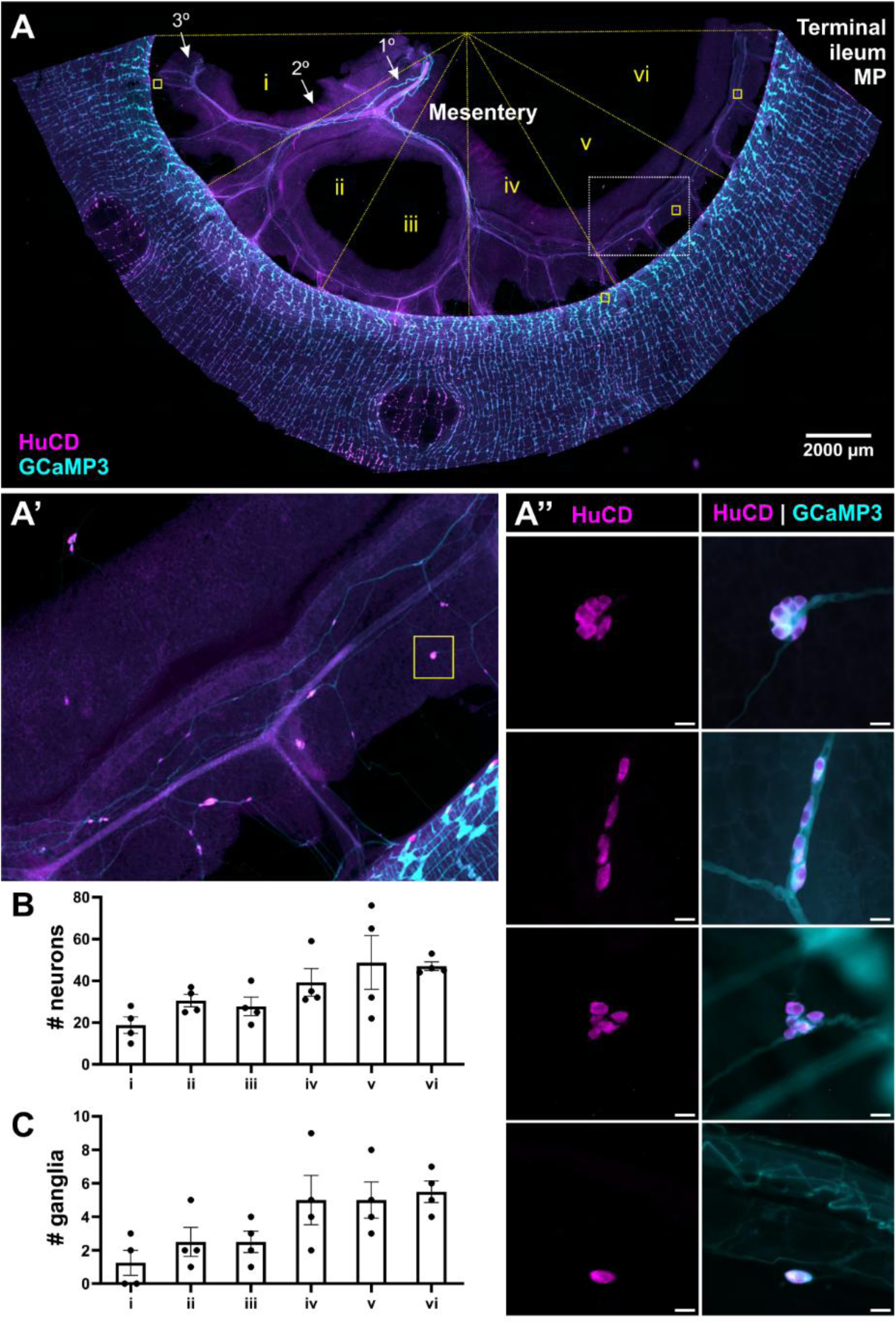
Neurons situated in the mesentery supplying the ileum. **A.** Representative tile scan of a flat sheet preparation of peeled terminal ileum myenteric plexus (MP) with the mesentery attached used for quantification of total neuronal and ganglion numbers (n = 4 preparations; N = 4 mice). The arrows indicate what we defined as the primary, secondary, and tertiary vascular branches. **A’.** Most neurons were situated around the tertiary branches and close to the gut wall as shown in the expanded view of the area marked by the white rectangle in **A**. Scale bar = 500 µm. **A”.** Higher magnification images of GCaMP3-expressing neurons (cyan) located in mesentery (marked by the yellow boxes in panel **A** labelled with the pan-neuronal marker HuCD (magenta). Scale bars = 20 µm. To quantify the distribution of neurons (**B**) and ganglia (**C**) along the length of the ileal mesentery, the mesentery was divided into six 30° sectors (**i**-**vi**), with the center point taken as the midpoint from the oral (left) to the anal end (right) of the ileal border as indicated by the yellow dotted lines in **A**. The total number of neurons and ganglia found associated with the tertiary vascular branches were counted within each sector. Only few neurons were found along the primary and secondary vascular branches, so these were excluded from quantification.

### Ganglia in the mesentery express typical enteric neuronal and glial markers

We also examined the neurochemistry of mesenteric neurons and glia using commonly used immunohistochemical markers for the ENS. First, mesenteric preparations were labelled with the neuronal markers HuCD and Phox2b, where 38/67 HuCD^+^ mesenteric neurons were Phox2b^+^ (N = 4 mice) (Tiveron et al., 1996;Lasrado et al., 2017). As in the ENS, ganglia in the mesentery displayed immunoreactivity not only for various neuronal markers, but also glial markers including sox10, glial fibrillary acidic protein (GFAP), and s100β (Boesmans et al., 2015) (N = 3 mice) (Figure 2A-C).

**Figure 2.**
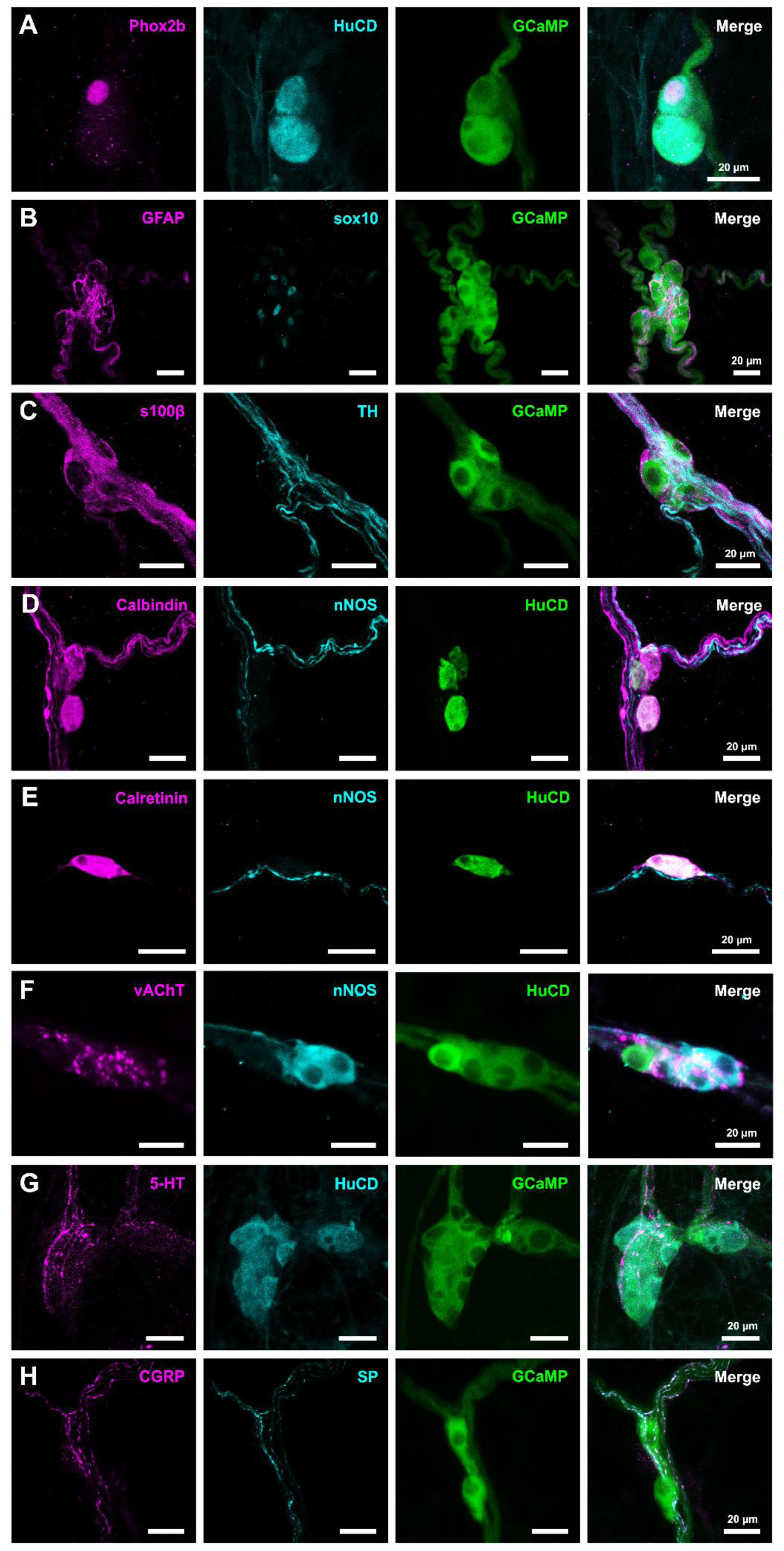
Neurons in the mesentery label for typical ENS markers. Representative confocal images of neurons and glia situated within mesenteric bundles supplying the ileum. Expression of the neuronal markers HuCD and Phox2b (**A**), as well as glial markers glial fibrillary acidic protein (GFAP), sox10, and s100β (**B-C**) were observed. Some mesenteric neurons labelled for (**D**) calbindin, (**E**) calretinin, or (**F**) neuronal nitric oxide synthase (nNOS). Varicosities immunoreactive for (**F**) vesicular acetylcholine transferase (vAChT), and (**G**) serotonin (5-HT) were seen in close apposition to mesenteric neurons. Mesenteric nerve fibers also labelled for (**H**) calcitonin gene-related peptide (CGRP), substance P (SP) and (**C**) tyrosine hydroxylase (TH). At least N = 3 mice were examined per marker.

The myenteric plexus largely comprises inhibitory nitrergic (nNOS^+^) neurons, and excitatory cholinergic calretinin^+^ and calbindin^+^ neurons (Qu et al., 2008). In the mesenteric bundles, 31/75 mesenteric neurons examined were nNOS^+^ (N = 6 mice), while 17/32 neurons were calbindin^+^ and 19/43 neurons were calretinin^+^ (N = 3 mice; Figure 2D-F). There was also some overlap in the expression of nNOS and calbindin (2/32 neurons examined), and nNOS and calretinin (5/38 neurons). In preparations stained for nNOS and calretinin, 7/38 neurons were negative for both markers, and in preparations labelled for nNOS and calbindin, 8/32 neurons were negative for both markers. Neurons in the mesentery were also found to be surrounded by cholinergic and serotonergic nerve varicosities, which labelled for vesicular acetyltransferase (vAChT) and 5-HT, respectively (N = 4 mice; Figure 2F-G). Serotonergic (i.e. 5-HT^+^) mesenteric neurons were rarely observed (1/38 neurons examined; N = 3 mice). Within the ENS, 5-HT^+^ neurons constitute only a small proportion (∼1%) of neurons in the mouse small intestine and are typically characterized as myenteric descending interneurons (Qu et al., 2008). Tyrosine hydroxylase (TH)-immunoreactivity was also observed and was most prominent in the mesenteric axon bundles but never in mesenteric neuronal cell bodies (0/17 neurons examined; N = 4 mice; Figure 2C). TH predominantly labels extrinsic sympathetic (noradrenergic) nerves, as well as a small proportion (<0.5%) of myenteric neurons (Qu et al., 2008). Additionally, varicosities immunoreactive for calcitonin gene-related peptide (CGRP) and substance P (SP) were also observed to closely appose mesenteric neurons (N = 3 mice) (Figure 2H). CGRP and SP marks spinal afferent fibers arising from the dorsal root ganglia (DRGs) (Qu et al., 2008; Sang & Young, 1996).

### Neurons in the mesentery respond to exogenous 5-HT, the nicotinic agonist DMPP, and ATP

To next examine whether mesenteric neurons are functionally active, calcium imaging was performed using Wnt1-Cre;R26R-GCaMP3 mice. Neurons and ganglia were identified based on their specific morphology and fluorescent GCaMP-expression. To determine whether mesenteric neurons respond to neurotransmitters or agonists commonly used to stimulate enteric neurons, serotonin (5-HT), DMPP (nicotinic agonist;) and ATP were applied by pressure ejection via a micropipette positioned adjacent to the neuron or ganglion of interest (Figure 3). Substances applied in this manner are expected to be diluted at least 1:10 when it reaches targeted cells (Breunig et al., 2007). 5-HT (1 mM) application evoked responses in 36/69 neurons examined (n = 19 preparations; N = 9 mice; Figure 3A-A”), while DMPP (100 µM) induced [Ca^2+^]_I_ transients in 53/67 neurons (n = 16 preparations; N = 9 mice; Figure 3B-B”). ATP (100 µM) elicited some neuronal as well as glial responses (21/69 neurons responded; n = 16 preparations; N=9 mice; Figure 3C-C”).

**Figure 3.**
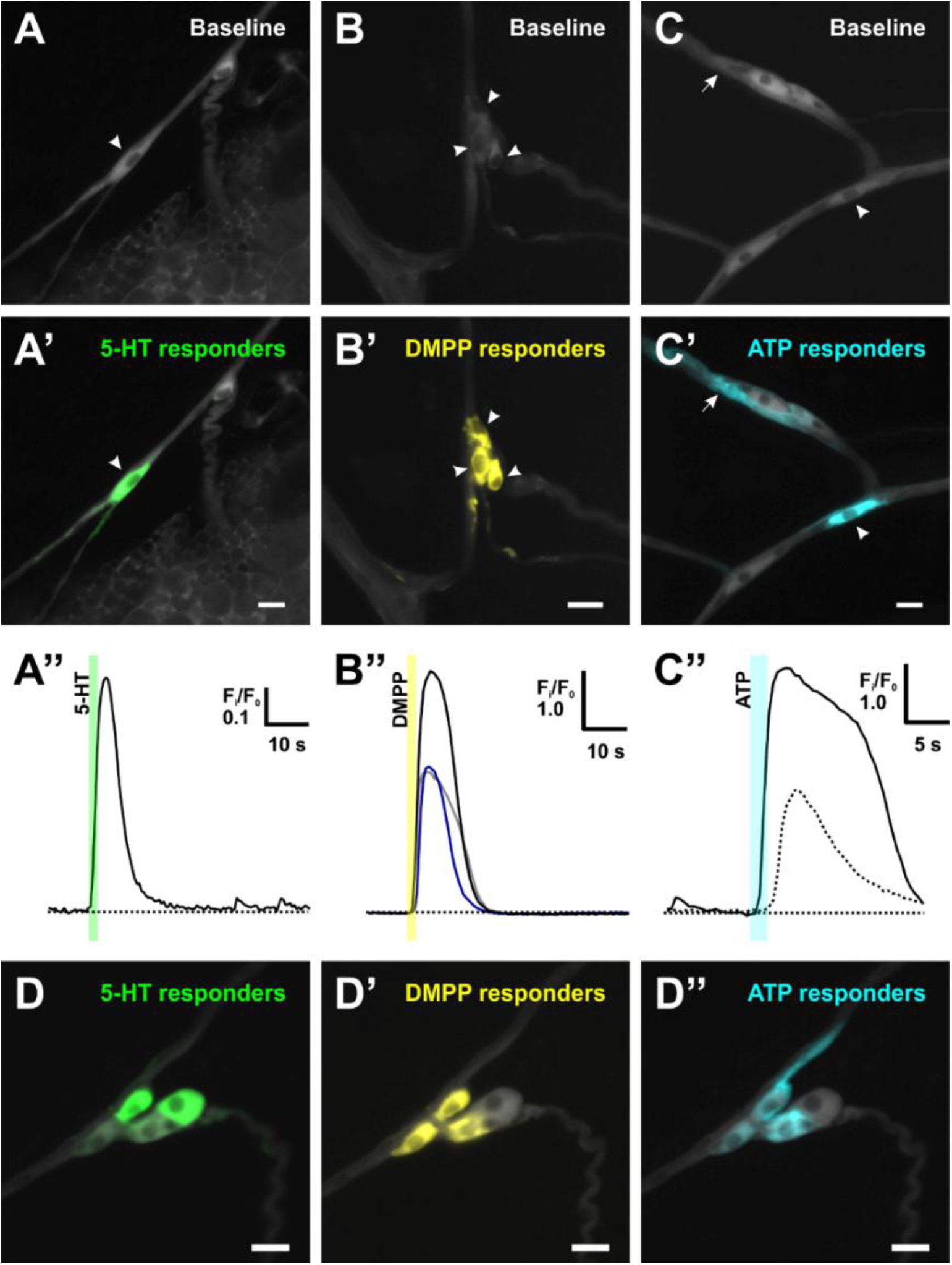
Mesenteric neurons respond to 5-HT, DMPP, and ATP. Representative widefield images of GCaMP3-expressing mesenteric neurons and glia situated within extrinsic nerve bundles supplying the ileum at baseline and their response to (**A-A”**) 5-HT (1 mM), (**B-B”**) DMPP (nicotinic agonist, 100 µM) and (**C-C”**) ATP (100 µM) applied by pressure ejection via a micropipette (10 psi; 2 s) at 10 s (N = 9 mice). Responding cells are highlighted using an overlay of a maximum minus average intensity projection (**A’-C’**). The corresponding traces depict the change in fluorescence over time (**A”-C”**) in the responding neurons and glia indicated by arrowheads or arrows, respectively. ATP also elicited a glial response (dotted line). In some preparations, the same neurons were sequentially exposed to (**D**) 5-HT, (**D’**) DMPP, and (**D”**) ATP. Some neurons responded to only one or multiple agonists applied (N = 3 mice). Scale bars = 20 µm.

In some experiments, 5-HT, DMPP and ATP were applied sequentially to the same ganglia, and the order of agonist application was alternated between preparations (N = 3 mice). Some neurons specifically responded only to 1 of the 3 agonists applied, while others responded to multiple or all 3 agonists (n = 3 preparations; N = 3 mice; Figure 3D-D”; Table 1). Of the three agonists tested, only ATP elicited glial responses.

**Table 1.**
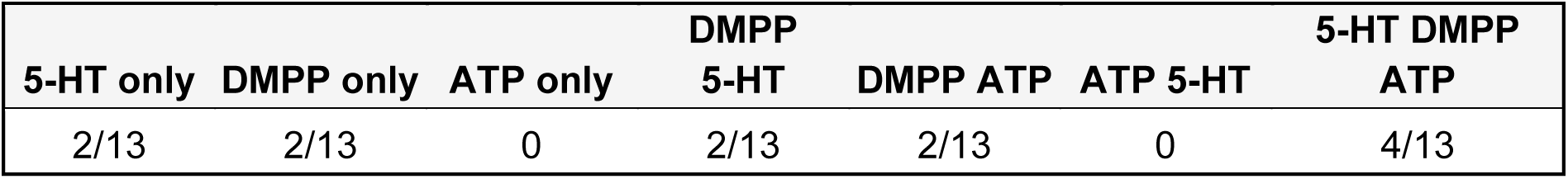
Number of mesenteric neurons that responded to 5-HT, DMPP and/or ATP locally applied via pressure ejection from a micropipette (n = 3 preparations; N = 3 mice).

### Myenteric neurons signal to mesenteric neurons via nicotinic transmission

Since mesenteric neurons were found to respond to transmitters and agonists used to excite the ENS (i.e. 5-HT, ATP, DMPP), we next examined whether they receive excitatory inputs from the ENS. To this end, we broadly depolarized the myenteric plexus and assessed mesenteric neuronal responses. For these experiments, we used ileum preparations that were cut open along the longitudinal axis approximately 1 mm from the mesenteric border such that the mesenteric connections remained intact on one side of the gut wall. The mucosa and submucosa of the ileum were then removed by microdissection to expose the myenteric and muscle layers. A sylgard divider was placed in the ring preparation along the mesenteric border and the bottom of the divider was sealed with silicone to create separate myenteric and mesenteric compartments as a means of confining the perfusion stimulus.

As intestinofugal neurons are typically found to be situated close to the mesenteric border (Furness et al., 2000; Tassicker et al., 1999; Timmermans et al., 1993), the stimulating pipette was positioned accordingly to target these neurons. High K^+^ (75mM) Krebs perfusion applied to the myenteric plexus and neuronal responses in the adjacent mesentery were examined (Figure 4). This elicited responses in 23/90 mesenteric neurons examined (32 preparations; N = 12 mice). The absence of responses in some neurons may be at least partly due to severed connections between the mesentery and the gut wall along the side that the ileum was cut open. Therefore, the number of mesenteric neurons that respond to myenteric activation is likely underrepresented. Spontaneous activity was observed in 6/32 neurons examined (N = 7 mice).

**Figure 4.**
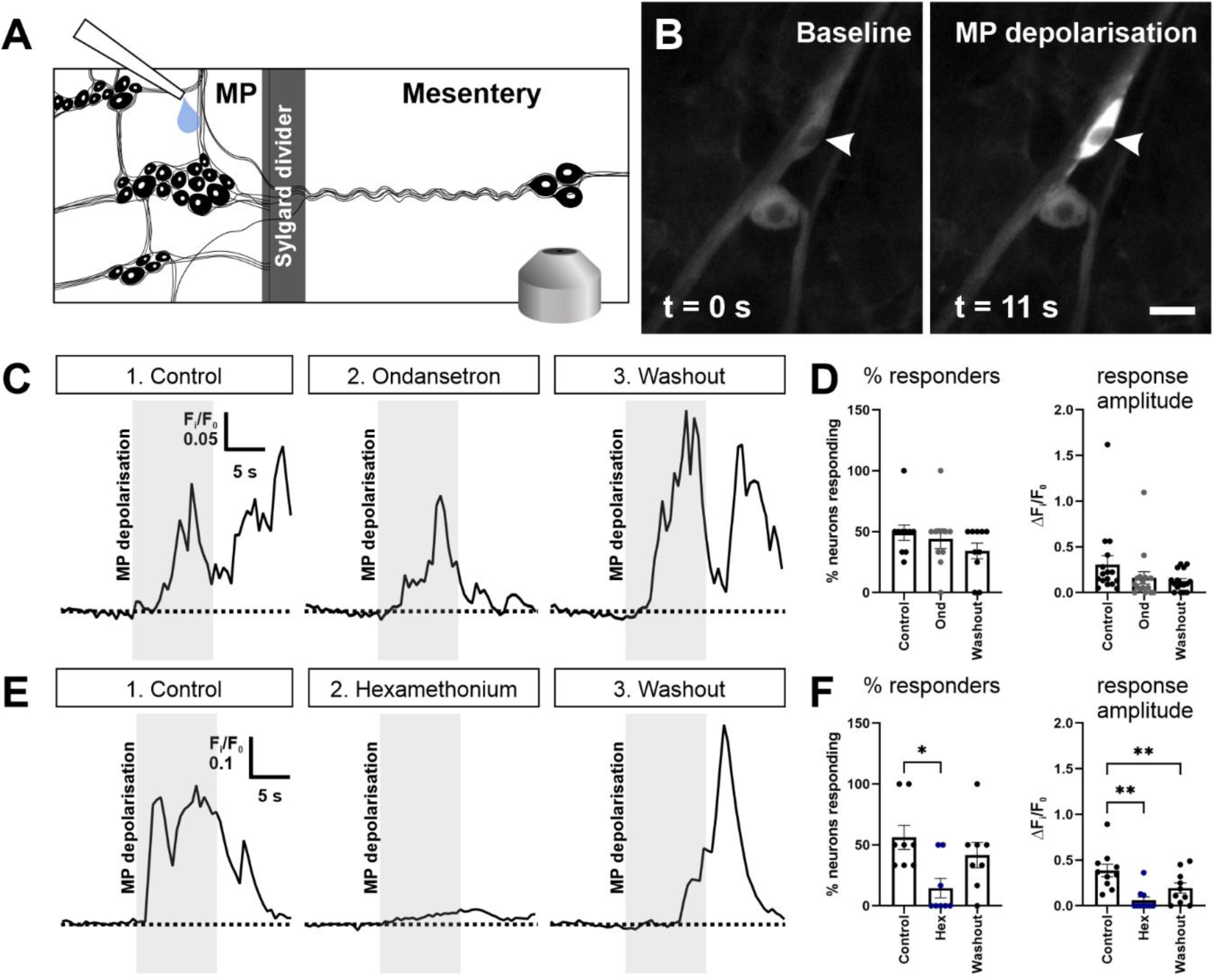
Mesenteric neurons respond to myenteric activation. Mesenteric neuronal responses to depolarisation of the myenteric plexus with high K^+^ perfusion (10 s) were assessed in flat sheet preparations consisting of peeled myenteric plexus with the mesentery attached on one side as shown in the schematic (**A**). The mesentery and myenteric plexus were separated by a sylgard divider positioned along the mesenteric border. The divider was sealed along the bottom edge with silicone. The perfusion tip was positioned close to the mesenteric border to target intestinofugal neurons. **B.** Representative images of a GCaMP3-expressing mesenteric neuron (indicated by arrowhead) that responded to depolarisation of the myenteric plexus under control conditions. **C.** Traces depict the change in fluorescence over time in a mesenteric neuron that responded to high K^+^ application to the myenteric plexus under 1) control conditions, 2) in the presence of a 5-HT_3_ antagonist ondansetron (ond; 10 µM; 10 min), and 3) after Krebs washout of the antagonist (N = 8 mice). Each stimulation was separated by 10 min. **D.** The neuronal response in the mesentery was not significantly affected in the presence of ondansetron. **E.** Traces depict the change in fluorescence over time in a mesenteric neuron that responded to high K^+^ application to the myenteric plexus under 1) control conditions, 2) during nicotinic blockade with hexamethonium (hex; 200 µM; 10 min), and 3) after Krebs washout of the antagonist (N = 4 mice). Each stimulation was separated by 10 min. **F.** The number of mesenteric neuronal responders to stimulation of the myenteric plexus, as well as their response amplitudes were significantly reduced by hex (Repeated measures one-way ANOVA, Dunnett’s multiple comparisons test; *p < 0.05, ** p < 0.01). This effect was partly reversed with washout.

As many mesenteric neurons responded to exogenously applied 5-HT and/or DMPP, we tested whether mesenteric neurons receive serotonergic or nicotinic excitatory inputs from the ENS. The 5-HT_3_ receptor antagonist ondansetron (10 µM) did not have a significant effect on mesenteric neuronal responses to myenteric high K^+^ application (Figure 5C, D) (27 mesenteric neurons examined; 10 preparations; N = 8 mice), indicating that serotonergic neurons do not contribute significantly to the mesenteric response observed in these preparations, at least not involving 5-HT_3_ receptors. One factor to consider is the length of the ileal preparation used. Previous studies in guinea pig ileum have demonstrated that serotonergic myenteric neurons are descending interneurons with long projections running distally (up to 24 mm) through the myenteric plexus (Furness & Costa, 1982). Therefore, our preparation (∼5 mm in length) may not necessarily encompass the full serotonergic neuron with its cell body in the myenteric plexus and its nerve terminal potentially synapsing onto a mesenteric neuron situated further distally. It is possible that the contribution of serotonergic signalling from the myenteric plexus to mesenteric neurons may only be apparent over longer distances.

**Figure 5.**
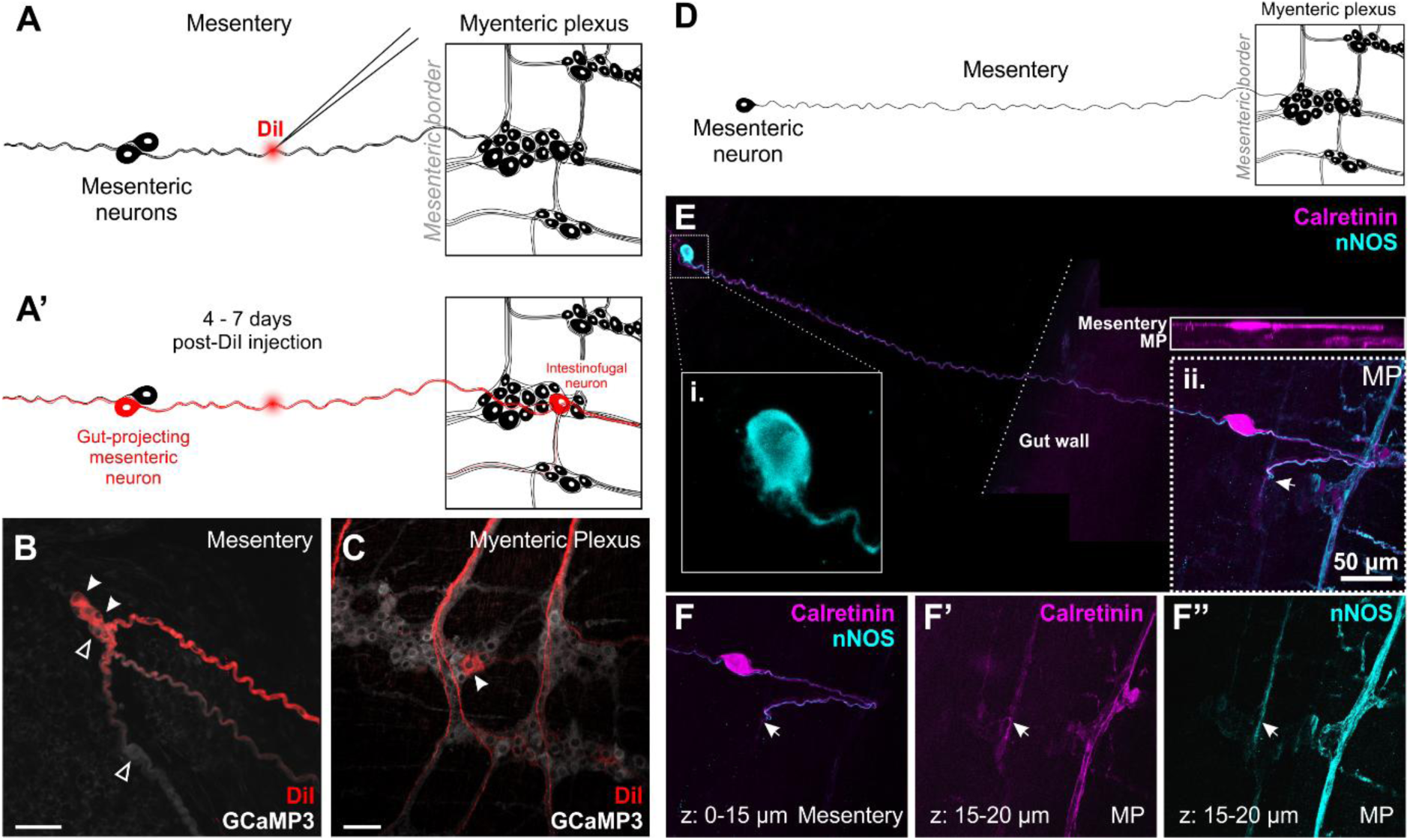
Mesenteric neurons project into the myenteric plexus. **A.** A schematic representation of a peeled myenteric plexus (MP) preparation used for DiI labelling, where the gut tube was opened ∼1-2mm to the side of the mesenteric border and mucosa and submucosa was removed by microdissection. DiI was locally applied via pressure ejection from a micropipette with its tip positioned in a nerve bundle in-between the gut wall and mesenteric neurons. **A’.** Preparations were assessed for DiI-labelling of neurons 4-7 days post-DiI injection. **B.** Some mesenteric neurons also show DiI labelling in some (arrowheads) but not all cell bodies (empty arrowheads). Thus, at least some mesenteric neurons have projections that run towards the gut wall (n = 6 preparations; N = 6 mice). **C.** A DiI labeled intestinofugual neuron found in the myenteric plexus situated close to the mesenteric border (n = 3 preparations; N = 3 mice examined). **D.** Schematic representation of the peeled ileum-mesentery preparation used for immunolabelling of calretinin and/or nNOS. In regions of mesentery where there were sparsely labelled nNOS^+^ and/or calretinin^+^ neurons, it was possible to follow the axon of these mesenteric neurons (n = 3/7 preparations; N = 3/7 mice) (**E-F**). **E.** Stitched confocal maximum projection images of an nNOS^+^ and a calretinin^+^ mesenteric neuron projecting into the myenteric plexus. **E** (**i**) The inset shows a single nNOS^+^ mesenteric neuron with an axon directed towards the gut wall. The dotted line marks the edge of the gut wall. The inset above the region indicated in (**ii**) shows the maximum projection in XZ, with a calretinin^+^ mesenteric neuron overlying the myenteric plexus. **F-F”** show maximum projections of sub-stacks of the region marked in **E** (**ii**), where **F** shows a calretinin^+^ mesenteric neuron as well as mesenteric fibers overlying the gut wall and **F’-F”** show the calretinin^+^ and nNOS^+^ mesenteric fibers leading into the myenteric plexus, respectively.

To next determine whether responses involve nicotinic transmission, mesenteric neuronal responses were assessed in the presence of hexamethonium (hex; 200 µM) (Figure 5E, F). Under control conditions, 56 ± 10% of mesenteric neurons responded (ΔFi/F0 = 0.39 ± 0.09; 10/23 neurons examined). Under hex conditions, responses were abolished in 7/10 neurons. Overall, the number of responders (15 ± 8% of neurons) and amplitude of responses were significantly reduced in the presence of hex (ΔFi/F0 = 0.06 ± 0.04) (n = 8 preparations; N = 4 mice; p = 0.01; repeated measures one-way ANOVA; Dunnet’s multiple comparisons test). This effect was partly reversed where 42 ± 10% of neurons responded (ΔFi/F0 = 0.19 ± 0.06; 8/23 neurons examined) following a 10 min Krebs washout.

### Anatomical connectivity between mesenteric neurons and the ENS

Given the proximity of mesenteric neurons to the gut wall and that they receive excitatory inputs from myenteric neurons, we were interested in next examining whether mesenteric neurons in turn signal back to myenteric neurons. To first address this anatomically, we assessed flat sheet preparations of peeled ileum myenteric plexus (with the mucosa and submucosa removed) and the mesentery attached and injected the lipophilic neuronal tracer DiI to nerve bundles emanating from the gut wall that were associated with mesenteric neurons (Figure 5A-A’). Some neuronal cell bodies in the mesentery were found to be labelled with DiI, providing additional evidence that some mesenteric neurons project into the gut wall (5/23 neurons, n = 6 preparations, N = 6 mice) (Figure 5B). Additionally, we observed some DiI-labelled myenteric intestinofugal neurons with their cell bodies situated close to the mesenteric border (Figure 5C) (n = 3 preparations; N = 3 mice) as described previously in rat, guinea pig, and pig (Furness et al., 2000; Tassicker et al., 1999; Timmermans et al., 1993).

Furthermore, in the same type of preparations that were immunolabelled for calretinin and nNOS (Figure 5D), we occasionally observed a single axonal projection arising from a calretinin^+^ or nNOS^+^ neuron in a sparsely labelled region in the mesentery (Figure 5D-F). Therefore, it was possible to follow the trajectory of these axons, and these were found to traverse into the myenteric plexus (in total 3 calretinin^+^ and 1 nNOS^+^ mesenteric neurons; n = 3/7 preparations examined; N = 7 mice). This indicates that at least some mesenteric neurons project into the ileum myenteric plexus.

### Mesenteric neurons signal to myenteric neurons

As some calretinin-IR mesenteric neurons were found to project into the myenteric plexus, we assessed whether mesenteric neurons provide excitatory inputs to myenteric neurons. However, given that only few neurons are present in the mesentery, mesenteric neuronal innervation of the myenteric plexus is likely sparse. Therefore, we used a lower magnification (10x) objective to image a larger field of view (1040 x 780 µm^2^) such that we could sample more myenteric neurons and increase the chance of finding a connection (if any). We stimulated mesenteric neurons using DMPP (100 µM) applied locally via a micropipette. When mesenteric neurons responded to DMPP (21/26 mesenteric neurons examined; n = 10 preparations; N = 4 mice), they were stimulated with DMPP at least four times (with a minimum of 3 min between each stimulation) to assess whether the mesenteric responses as well as any myenteric responses were reproducible (Figure 6). Myenteric responses were observed in response to DMPP stimulation of 6/26 mesenteric neurons, and myenteric responses were consistently elicited in 4/10 preparations examined. These data are comparable to the proportion of DiI-labelled mesenteric neurons found projecting towards the gut (5/23 neurons examined). Of these preparations where myenteric responses were observed, up to 4 myenteric neurons per preparation displayed consistent responses to mesenteric DMPP stimulation (in total, 7/580 myenteric neurons examined responded). While only a small proportion of myenteric neurons responded, this is likely an under-representation due to two key considerations. Firstly, we severed at least half of the connections running between the mesentery and the gut wall to make the flat sheet preparations necessary for imaging. Secondly, we have no prior knowledge of where the mesenteric neurons stimulated send their axons and where their nerve terminals end within the myenteric plexus. Therefore, we may have missed potential responses beyond the field of view. Finally, another possibility is that mesenteric neurons also provide inhibitory, rather than excitatory, inputs to the myenteric plexus considering some mesenteric neurons are nNOS^+^.

**Figure 6.**
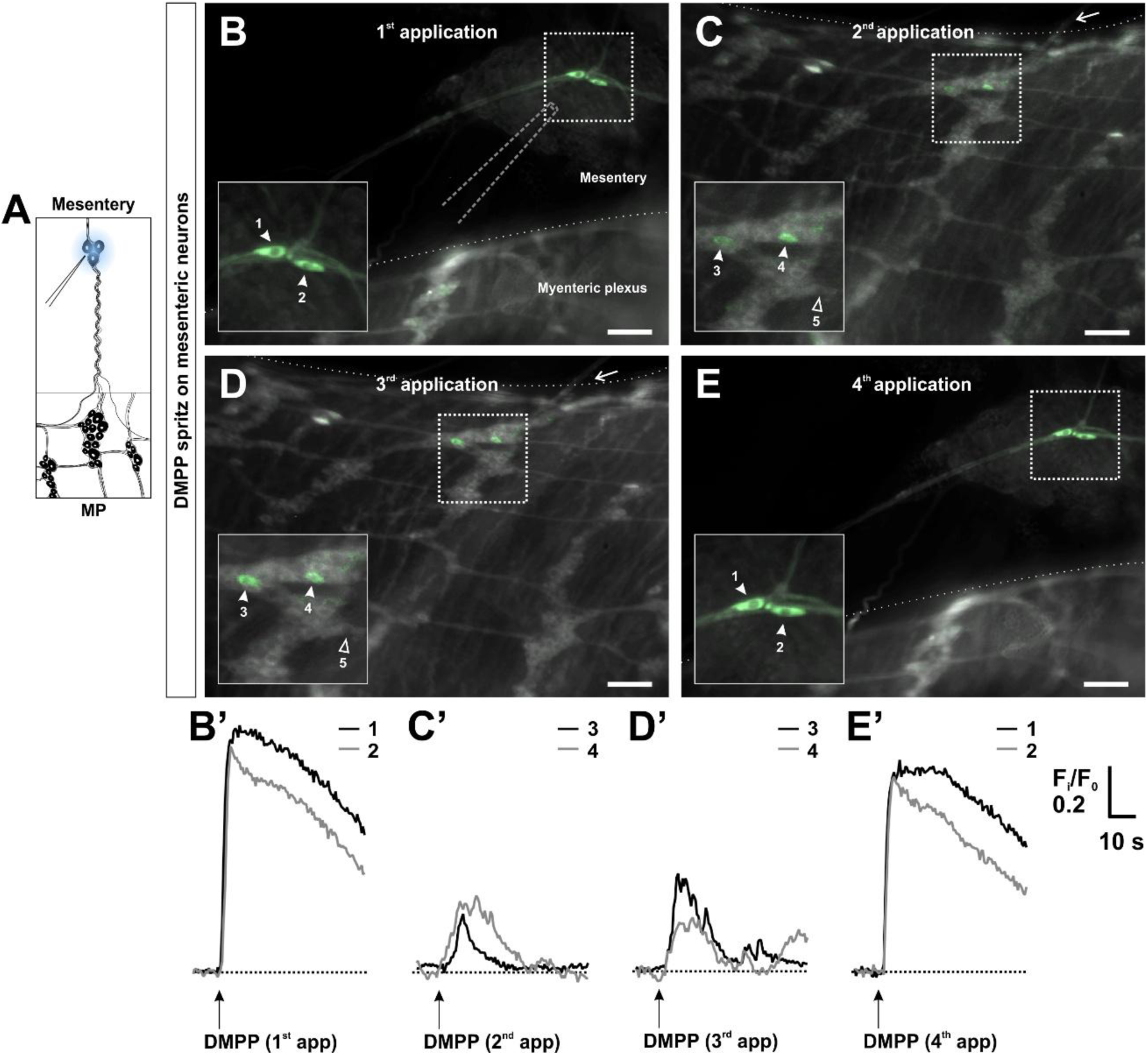
Myenteric neurons receive excitatory inputs from mesenteric neurons. **A.** DMPP (100 µM) was locally applied to the mesenteric neurons via a micropipette. **B-E’.** DMPP was applied to the mesenteric neurons consecutively 4 times (n = 10 preparations; N = 4 mice). The dotted line demarcates the mesenteric border. The arrow indicates a nerve bundle running between the targeted mesenteric neurons and the myenteric plexus Consistent responses were elicited in both the mesenteric neurons themselves (**B-B’**, **E-E’**) and also in some myenteric neurons close to the mesenteric border (**C-C’**, **D-D’**). Responding cells are highlighted using a maximum minus average intensity projection (green) overlay on an average intensity projection (greyscale). Scale bars = 100 µm.

### No evidence for any association between mesenteric neurons and other extrinsic targets

The sympathetic innervation of the small intestine is provided by neurons with their cell body situated in the celiac-superior mesenteric ganglion complex (CG-SMG) and their axon runs through the mesentery into the gut wall (Muller et al., 2020; Szurszewski & Linden, 2012). Intestinofugal neurons in the ileum in turn project out of the gut wall and through the mesentery to the CG-SMG (Muller et al., 2020; Szurszewski & Linden, 2012). Considering the location of the neurons found along mesenteric nerve fiber tracts, we examined whether they may also receive signals from or send signals to neurons in the CG-SMG. In peeled ileal MP preparations with the CG-SMG and associated mesenteric connections attached (Figure 8A), we applied an electrical train stimulus (20 Hz, 2 s) using a focal electrode positioned approximately on the medial part of the SMG (Wang et al., 2025) and assessed for responses in mesenteric neurons. No responses were observed in the cell bodies of mesenteric neurons (n = 24 mesenteric neurons examined; N = 2 mice). In some instances, the mesenteric neurons were situated along nerve fiber tracts that displayed a Ca^2+^ response to the electrical stimulus, but the cell bodies themselves did not respond (10/24 mesenteric neurons examined; N = 2/3 mice; Figure 7C). In addition, responses were observed in myenteric neurons in the ileum indicating that there were intact extrinsic nerve connections running through the mesentery to the gut in these preparations (N = 2/3 mice; Figure 7D).

**Figure 7.**
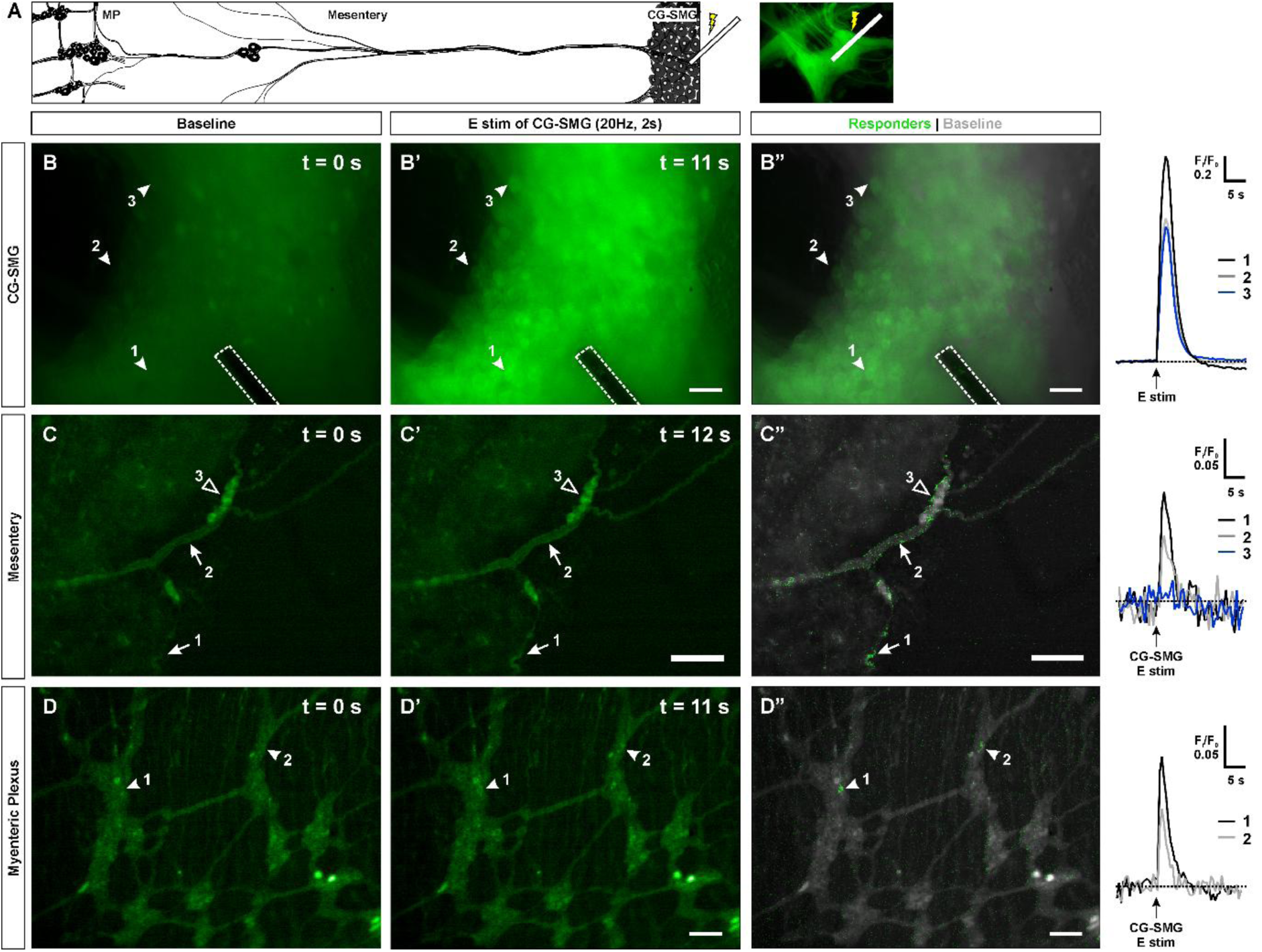
CG-SMG stimulation elicits responses in the myenteric plexus but not in mesenteric neurons. **A.** Schematic showing the type of preparation used for Ca^2+^ imaging, which comprised the celiac-superior mesenteric ganglion complex (CG-SMG) with peeled ileal myenteric plexus and the associated mesenteric connections intact. A focal electrode was positioned on the CG-SMG to apply an electrical stimulus. **B-B”** Representative images of the CG-SMG at baseline and during focal electrical stimulation (20 Hz, 2s). The dotted line outlines the focal electrode positioned on the ganglion. Example traces of responding postganglionic sympathetic neurons are indicated by corresponding numbered arrowheads in panels B-B”. The pulse-train stimulus was applied at 10 s as shown by the arrow. **C-C”**. Electrical stimulation of the CG-SMG evoked Ca^2+^ transients in nerve fibers (arrows) but not the neuronal cell bodies (empty arrowhead) in the mesentery (example traces shown in the right panel). **D-D”.** Activating the CG-SMG also elicited responses in the myenteric plexus (indicated with arrowheads and traces depicted in the right panel) (n = 2 preparations; N = 2 mice). Scale bars = 50 µm.

**Figure 8.**
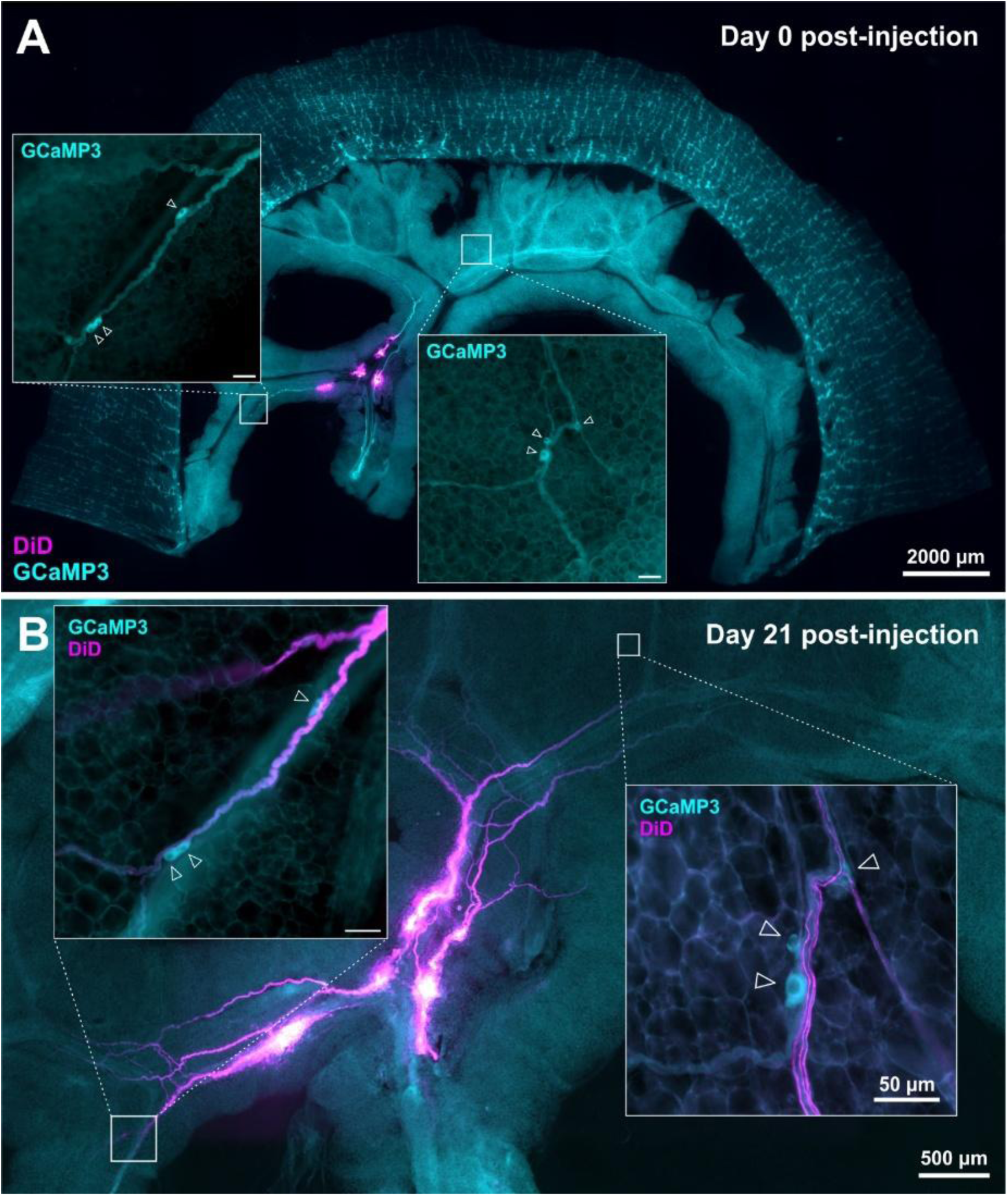
DiD tracing from mesenteric nerve trunks do not label mesenteric neurons. **A.** In Wnt1xGCaMP3 peeled terminal ileum preparations with associated mesentery attached, DiD (magenta) was applied onto exposed severed mesenteric nerve trunks. Insets show some mesenteric neurons found along the mesenteric nerve bundles at Day 0 post-injection. **B.** Mesenteric neurons (cyan) were assessed for labelling 21 days post-DiD application. While DiD labelling of mesenteric nerve fiber bundles (magenta) was extensive, none of the GCaMP^+^ neuronal cell bodies (cyan; indicated by arrowheads) found in the mesentery were DiD^+^ (n = 3 preparations; N = 3 mice). Scale bars of inset panels = 50 µm.

Further, we injected DiD into mesenteric nerve trunks severed from the spinal cord side to more generally assess whether mesenteric neurons have projections towards other extrinsic neuronal targets including the nodose, dorsal root, or prevertebral sympathetic ganglia. DiD-labelled preparations were imaged at least 7 days and up to 28 days post-DiD injection. While we readily identified DiD labelling of extrinsic nerve fibers running throughout the mesentery, we did not observe any labelling of mesenteric neuronal cell bodies situated alongside the DiD^+^ fibers (Figure 8; n = 0/45 neurons examined; N = 3 preparations).

### Mesenteric neurons respond to ileal distension

Since mesenteric neurons appear to preferentially interact with the ENS, we further investigated what functional role they may play in intestinal physiology. A previous study on the nerve of Remak in chicks showed that these extra-intestinal neurons are responsive to intestinal distension and the authors speculated that they may have a role in regulating gut motility (Smith & Lunam, 1998). Therefore, we explored the possibility that mesenteric neurons may have a similar function. We cannulated a segment of terminal ileum (∼5 cm) with the mesentery attached and then injected Krebs into the intestinal lumen as a means of distending the gut, while performing Ca^2+^ imaging of activity in mesenteric neurons (Figure 9). Indeed, some mesenteric neurons consistently responded to repeated distension of the ileum (10/15 neurons examined; N = 4 mice).

**Figure 9.**
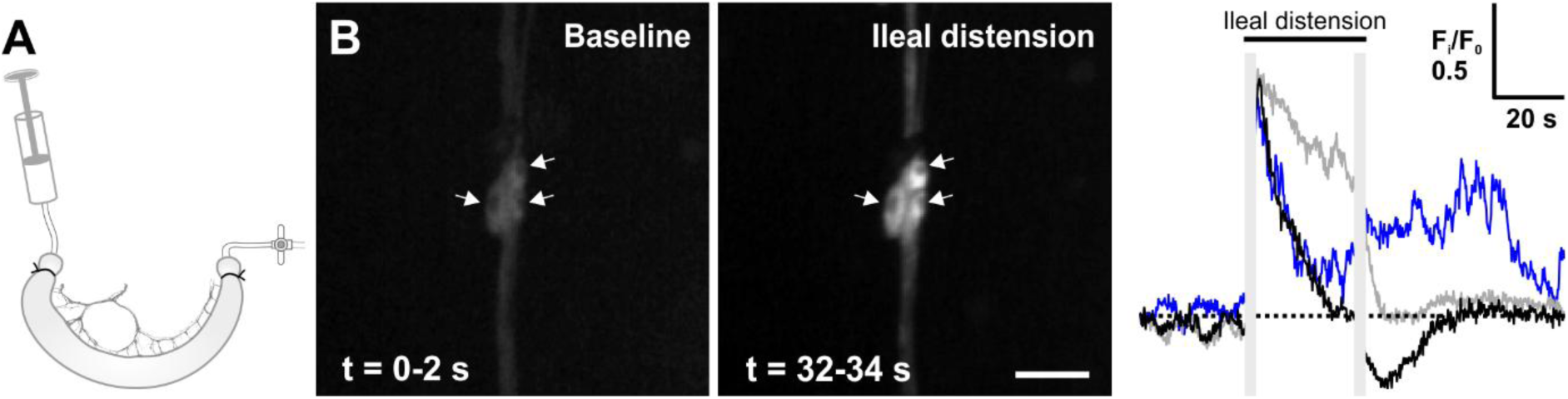
Mesenteric neurons respond to ileal distension. **A.** Schematic of the preparation used for imaging. Using a fluorescence stereomicroscope, Ca^2+^ activity was imaged in neurons found in the mesentery connected to a segment of cannulated terminal ileum from Wnt1xGCaMP3 mice. We assessed for mesenteric neuronal responses to ileal distension evoked by injecting ∼300 µl of Krebs solution into the lumen (N = 4 mice). **B.** Average projections of the fluorescence at baseline and during ileal distension. Scale bar = 50 µm. Traces of the Ca^2+^ transients in responding mesenteric neurons indicated by arrows. The signal in frames with motion artefacts immediately before and after distension were removed (replaced by grey bars) as the neurons shifted out of focus due to the gut distension.

### Mesenteric neurons provide inhibitory modulation of ENS signaling around the circumference of the gut

Having found that mesenteric neurons are responsive to intestinal distension, we next considered whether they provide feedback to the enteric network. Given the location of mesenteric neurons and their projections into the gut wall, we examined whether they contribute to signaling in pathways within the myenteric plexus, particularly circumferentially across the mesenteric border. We used peeled myenteric plexus while leaving the mesentery attached by opening the gut along the anti-mesenteric border. Myenteric responses were evoked using local 5-HT (100 µM) application by pressure ejection onto a ganglion on one side of the mesenteric border. Responses were first assessed at the site of stimulation and then in ganglia situated circumferentially across the mesenteric border. We next repeated this set of stimulations in the same preparation after dissecting away the mesentery. In control experiments, two sets of stimulations were performed using the same preparation with the mesentery intact throughout. The two locations imaged either side of the mesentery were typically ∼2 ganglia apart in the circumferential direction (controls: 1.44 ± 0.06 mm, n = 9 paired locations imaged; N = 5 mice vs. with and without mesentery: 1.63 ± 0.12 mm; n = 10 paired locations imaged; N = 4 mice) (Figure 10A, B). Up to 4 paired locations were assessed per preparation.

**Figure 10.**
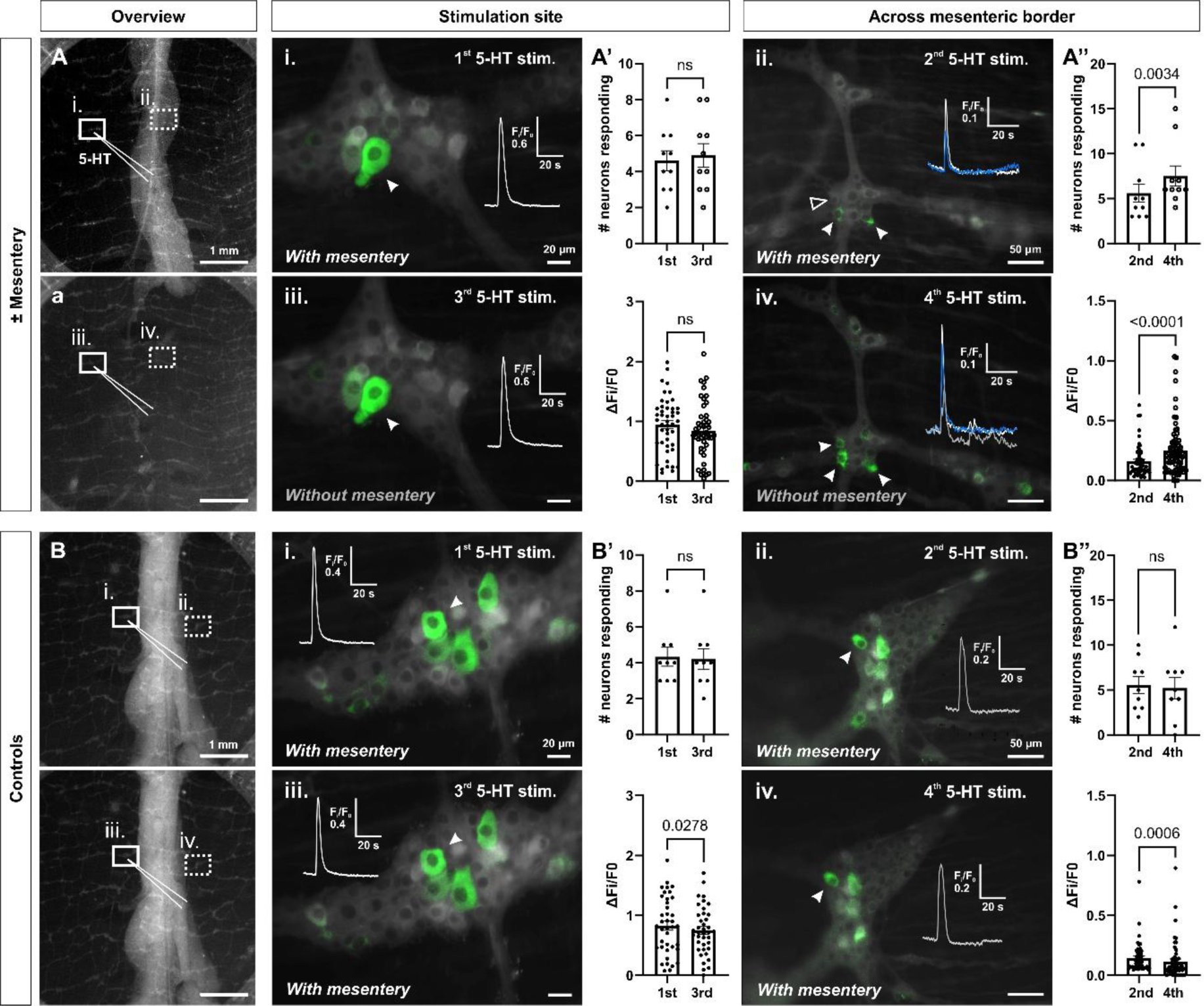
The mesentery provides a source of inhibition to the myenteric circuitry. Using Wnt1xGCaMP3 mice, Ca^2+^ imaging was performed on preparations of peeled myenteric plexus with the mesentery attached. **A.** 5-HT (100 µM) was applied twice to a selected ganglion (**i**) by pressure ejection via a micropipette (10 psi; 2 s) at 10 s. Each application was separated by at least 3 min. Responses were imaged at the stimulus site and then (**ii**) in a field of view (464 x 351 µm^2^) located circumferentially across the mesenteric border (∼2 ganglia apart). Responding cells are highlighted in green using an overlay of the maximum minus average intensity projection over the baseline fluorescence (grayscale). Examples of responding neurons are indicated by arrowheads and the corresponding traces depicting the change in fluorescence over time are shown in the inset. The set of 5-HT stimulations were repeated in the same preparation after dissecting away the mesentery (**a**, **iii**, **iv**). The upper panels of graphs show the number of responding neurons and the lower panels show the 5-HT response amplitudes at (**A’**) the stimulation site and (**A”**) across the mesenteric border from the stimulation site. Across the mesenteric border, both the number of responders and response amplitudes increased after removing the mesentery (paired t-test; n = 10 fields of view either side of the mesenteric border imaged; N = 4 mice). **B-B”.** In control experiments, the set of 5-HT stimulations were repeated, and responses were assessed either side of the mesenteric border as described in **A** but the mesentery remained intact throughout. Response amplitudes at the stimulation site and across the mesenteric border were significantly reduced with repeated 5-HT stimulation (paired t-test; n = 9 fields of view either side of the mesenteric border imaged; N = 5 mice).

First, the number of neurons that responded to 5-HT at the stimulus site was similar between controls, and preparations with and without mesentery (4 ± 0.5 vs. 5 ± 0.6 neurons; n = 9-10 ganglia examined, respectively) (Figure 10). Response amplitudes were also similar with mesentery: ΔFi/F0 = 0.96 ± 0.07 and without mesentery: ΔFi/F0 = 0.85 ± 0.07 (Figure 10A’). However, response amplitudes were significantly reduced in controls with repeated 5-HT application (1^st^: ΔFi/F0 = 0.83 ± 0.08 vs. 3^rd^: ΔFi/F0 = 0.75 ± 0.06; paired t-test: p = 0.0278) (Figure 10B’). On the other hand, response amplitudes of neurons in ganglia located circumferential to the stimulus site significantly increased after the mesentery was removed (ΔFi/F0 = 0.16 ± 0.02 vs. ΔFi/F0 = 0.25 ± 0.02; p < 0001) (Figure 10A”). By contrast, corresponding response amplitudes in controls were significantly decreased (ΔFi/F0 = 0.14 ± 0.02 vs. ΔFi/F0 = 0.12 ± 0.02; paired t-test: p = 0006) (Figure 10B”). Even though there was no significant difference in the number of neurons that responded at the stimulation site with or without mesentery, the number of responders increased across the mesenteric border after removing the mesentery (with mesentery: 18 ± 3% vs. without mesentery: 25 ± 4% neurons examined; p = 0.0034) (Figure 10A’-A”). This was not observed in control preparations (16 ± 2% vs. 15 ± 3% neurons examined) (Figure 10B’-B”). Taken together, these data indicate that the mesentery can provide a source of inhibition to the myenteric plexus.

## Discussion

In this study, we describe a population of extrinsic neurons found in the mesentery which supply and interact with the ENS (Figure 11). These mesenteric neurons form a neuro-glial network that is reminiscent of the submucosal plexus and runs alongside the small intestinal tube. In terms of their neurochemistry, mesenteric neurons share common markers with enteric neurons such as nNOS, calbindin, and calretinin, but do not express TH which is a marker of sympathetic nerves. With DiI and DiD neuronal tracing as well as calcium imaging, mesenteric neurons were found to be both anatomically and functionally connected to neurons in the myenteric plexus of the ileum and there is bidirectional communication between these two populations of neurons.

**Figure 11.**
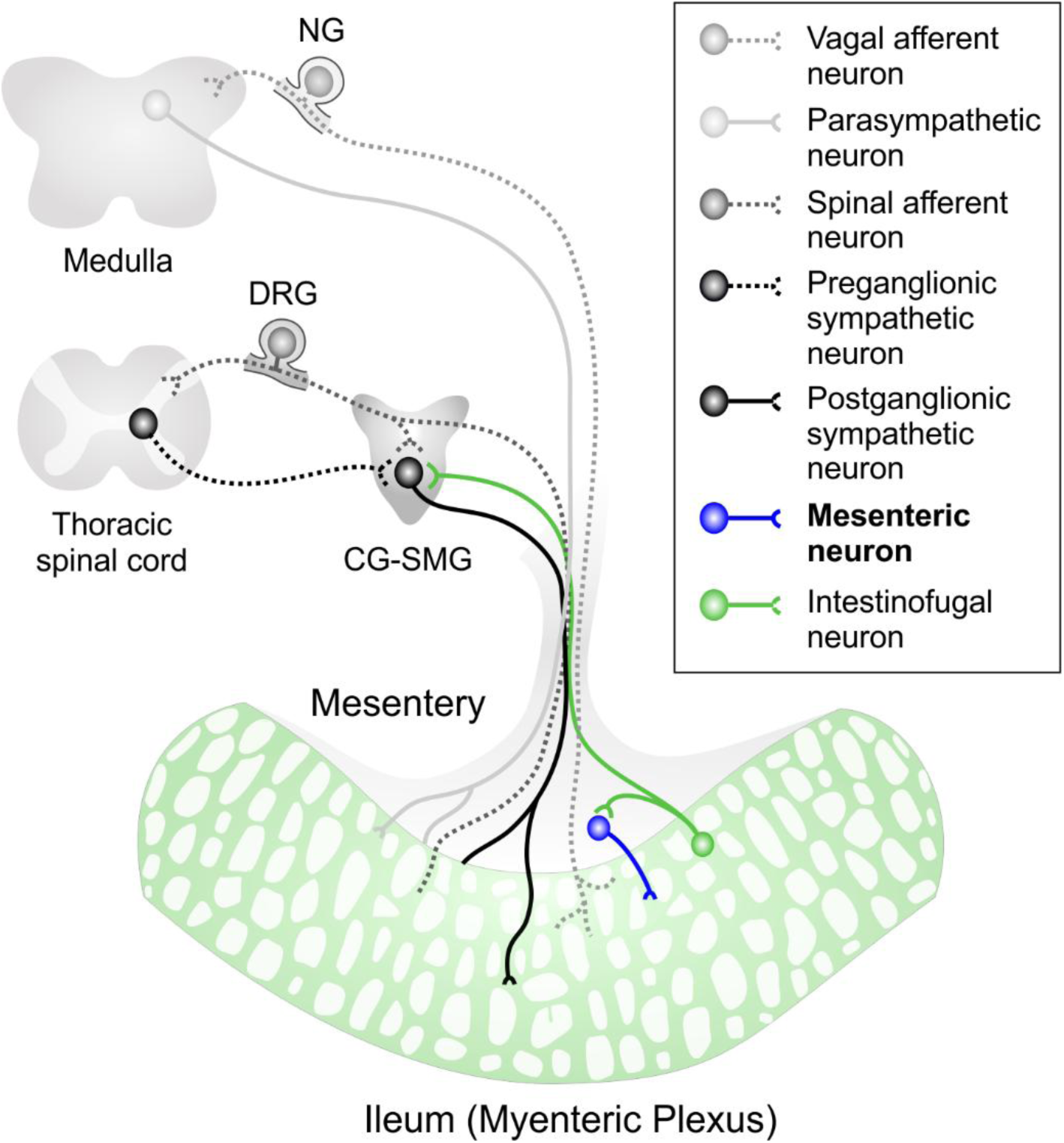
Summary schematic of the various sources of extrinsic innervation supplying the small intestinal myenteric plexus. These include vagal afferents from the nodose ganglion (NG), spinal afferents from, the dorsal root ganglia (DRGs), preganglionic parasympathetic neurons from the dorsal motor nucleus in the brainstem, postganglionic sympathetic neurons from the prevertebral ganglia (celiac- and superior mesenteric ganglion complex; CG-SMG), as well as mesenteric neurons. Mesenteric neurons in turn receive excitatory (nicotinic) inputs from myenteric intestinofugal neurons.

Considering the location of mesenteric neurons, we also investigated whether they interact with other (extrinsic) neuronal targets, but this does not appear to be the case. Previous work suggests that prevertebral sympathetic ganglia and intestinofugal neurons arising from the gut wall are involved in relaying long-distance physiological or pathophysiological signals from distal to more proximal regions of the GI tract (Stebbing et al., 2024; Zhang et al., 2022). The axonal projections of these postganglionic neurons and intestinofugal neurons run through the mesentery. However, we did not find evidence that mesenteric neurons (some of which sit along these extrinsic nerve bundles) communicate with the CG-SMG, nor do they project to other extrinsic neuronal targets. Thus, our data indicate that mesenteric neurons preferentially interact with the ENS. In addition to this functional connection between mesenteric and enteric neurons, whether there is also a developmental association between these two neuronal populations needs to be further explored.

Finally, we found that local intestinal distension can activate mesenteric neurons and that mesenteric neurons in turn provide inhibitory inputs to circumferentially directed pathways activated by local 5-HT application. 5-HT can activate intrinsic primary afferent neurons (IPANs) (Fung et al., 2025), which are mechanosensitive (Bornstein, 2006; Mao et al., 2006) and form recurrent networks primarily along the circumferential axis (Stebbing & Bornstein, 1994; Thomas et al., 2000). Overall, these data indicate that mesenteric neurons can provide inhibitory inputs as a form of negative feedback to the gut and potentially limit the level of excitation within the myenteric neurocircuitry i.e. in the event of a distension stimulus. Thus, mesenteric neurons can contribute to modulating signaling within the enteric neural network and may be important for maintaining network stability (Thomas & Bornstein, 2003). Future experiments comparing ileal motility with and without the mesentery will be important to further extend our understanding of the contribution of mesenteric neurons to regulating intestinal motility in health and in disease.

## Materials and Methods

### Animals

Adult mice from at least 6 weeks of age and of either sex were used including *Wnt1-Cre;R26R-GCaMP3* mice in which the genetically-encoded Ca^2+^ indicator GCaMP3 is expressed in all neural crest-derived cells. For calcium imaging experiments, *Wnt1-Cre;R26R-GCaMP3* mice were used. All mice were sacrificed by cervical dislocation. All procedures were approved by the animal ethics committee of the University of Leuven (KU Leuven) guidelines for the use and care of animals.

### Quantification of neurons and ganglia in the ileal mesentery

Immunohistochemical stainings were visualised using a Zeiss Axio Imager M2 with an Axiocam 705 (Zeiss). Each preparation was completely screened for mesenteric neurons by imaging both sides of the tissue (i.e. first with the circular muscle side up and then the serosal side or vice versa) as the layer of mesenteric adipose tissue can obscure some of the neurons from one side. The numbers of neurons and ganglia per field of view (675 x 563 µm^2^) were counted using a 20x objective and an overview image of the entire preparation was made by performing a tile scan using a 5x objective lens.

### Immunohistochemistry

To assess the numbers and neurochemical coding of the extrinsic neurons and ganglia in the ileal mesentery, immunohistochemical labeling was performed. The mesentery bundle preparations and Ca^2+^ imaging preparations were fixed for 2 h in 4% paraformaldehyde (PFA, Merck) in 0.1 M phosphate buffered saline (PBS, pH = 7.3-7.4) and were then washed in PBS (3 × 10 min). These preparations were incubated with a blocking buffer containing 4% donkey and/or goat serum (Jackson Immuno Labs) and 2% triton X-100 (Sigma Aldrich) in PBS for 2 h at room temperature, and then were incubated with various combinations of primary antisera (Table 2) at least overnight and up to 72 h at 4°C. Following PBS washes (3 × 10 min), preparations were incubated with secondary antisera (Table 3) for 2 h at room temperature. Primary and secondary antisera applied were diluted in blocking buffer. Secondary antisera were applied sequentially in instances where combinations of primary and secondary antisera were both raised in goat or sheep. Preparations were washed in PBS (3 x 10 min) prior to mounting onto slides using Citifluor (Citifluor Ltd., Leicester, UK). Preparations were imaged on either a Zeiss LSM 780, LSM 880, or LSM 980 laser scanning confocal microscope, and/or a Zeiss AxioImager widefield microscope.

**TABLE 2.**
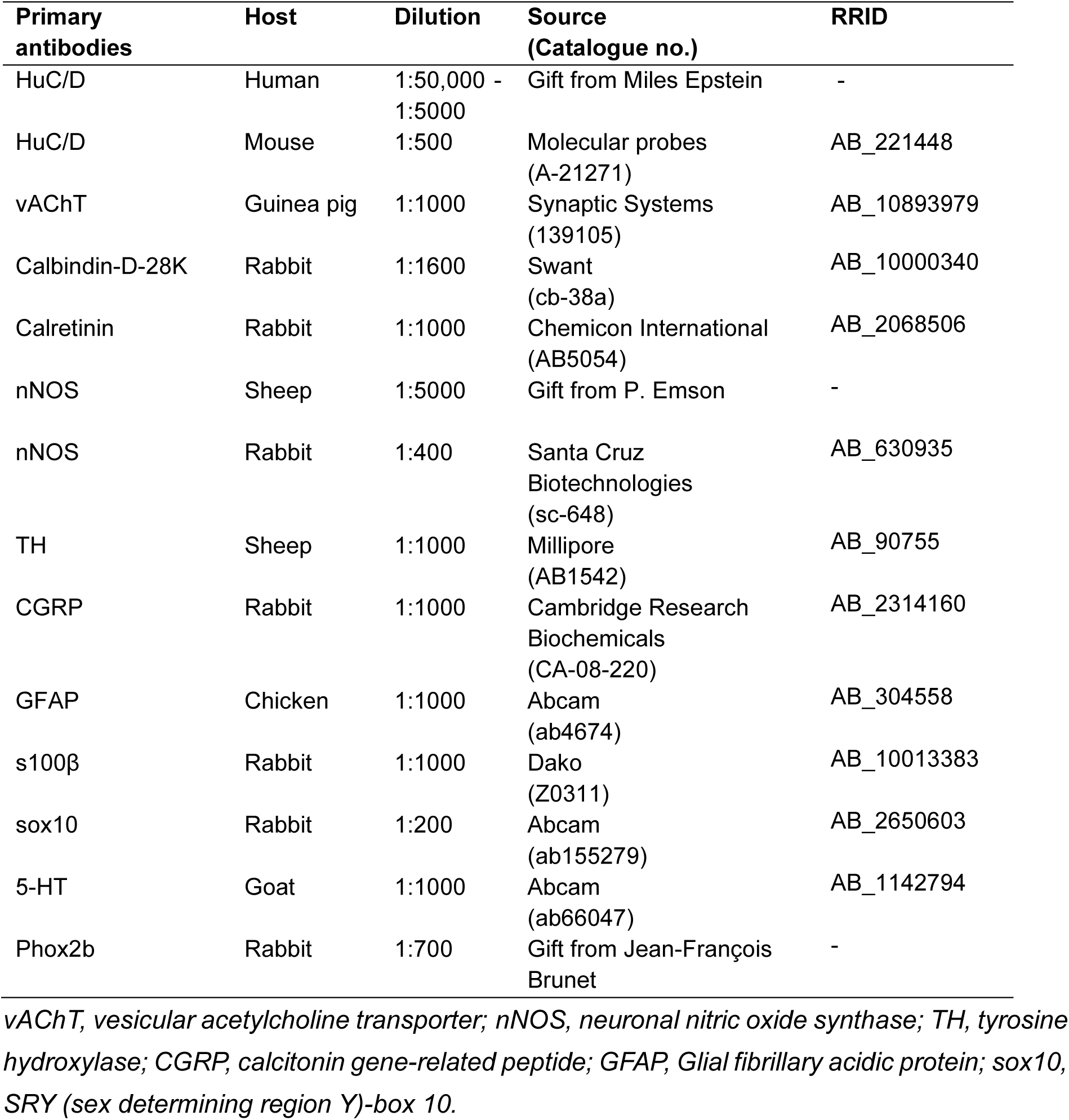
Primary antibodies used for immunohistochemistry.

| Primary antibodies | Host | Dilution | Source (Catalogue no.) | RRID |
| --- | --- | --- | --- | --- |
| HuC/D | Human | 1:50,000 - 1:5000 | Gift from Miles Epstein | - |
| HuC/D | Mouse | 1:500 | Molecular probes (A-21271) | AB_221448 |
| vAChT | Guinea pig | 1:1000 | Synaptic Systems (139105) | AB_10893979 |
| Calbindin-D-28K | Rabbit | 1:1600 | Swant (cb-38a) | AB_10000340 |
| Calretinin | Rabbit | 1:1000 | Chemicon International (AB5054) | AB_2068506 |
| nNOS | Sheep | 1:5000 | Gift from P. Emson | - |
| nNOS | Rabbit | 1:400 | Santa Cruz Biotechnologies (sc-648) | AB_630935 |
| TH | Sheep | 1:1000 | Millipore (AB1542) | AB_90755 |
| CGRP | Rabbit | 1:1000 | Cambridge Research Biochemicals (CA-08-220) | AB_2314160 |
| GFAP | Chicken | 1:1000 | Abcam (ab4674) | AB_304558 |
| s100β | Rabbit | 1:1000 | Dako (Z0311) | AB_10013383 |
| sox10 | Rabbit | 1:200 | Abcam (ab155279) | AB_2650603 |
| 5-HT | Goat | 1:1000 | Abcam (ab66047) | AB_1142794 |
| Phox2b | Rabbit | 1:700 | Gift from Jean-François Brunet | - |

*vAChT*, vesicular acetylcholine transporter; *nNOS*, neuronal nitric oxide synthase; *TH*, tyrosine hydroxylase; *CGRP*, calcitonin gene-related peptide; *GFAP*, Glial fibrillary acidic protein; *sox10*, *SRY* (sex determining region Y)-box 10.

**TABLE 3.** Secondary antibodies used for immunohistochemistry.

| Secondary antibodies | Host | Dilution | Source (Catalogue no.) | RRID |
| --- | --- | --- | --- | --- |
| Anti-Human AMCA | Goat | 1:250 | Jackson ImmunoResearch (109-155-003) | AB_2337696 |
| Anti-Guinea pig AF594 | Goat | 1:1000 | Molecular Probes (A-11076) | AB_141930 |
| Anti-Rabbit AF647 | Donkey | 1:500 | Molecular Probes (A-31573) | AB_2536183 |
| Anti-Rabbit AF594 | Donkey | 1:250-1:1000 | Molecular Probes (A-21207) | AB_141637 |
| Anti-Chicken AF594 | Goat | 1:1000 | Molecular Probes (A-11042) | AB_2534099 |
| Anti-Goat Cy3 | Donkey | 1:500 | Jackson Immuno Labs (705-165-147) | AB_2307351 |
| Anti-Rabbit AMCA | Donkey | 1:250 | Jackson ImmunoResearch (711-155-152) | AB_2340602 |
| Anti-Human AF594 | Donkey | 1:1000 | Jackson ImmunoResearch (709-585-149) | AB_2340572 |
| Anti-Goat AF647 | Donkey | 1:1000 | Thermo Fisher Scientific (A-21447) | AB_2535864 |
| Anti-Sheep AF647 | Donkey | 1:1000 | Thermo Fisher Scientific (A-21448) | AB_2535865 |
| Anti-Mouse AMCA | Donkey | 1:250 | Jackson ImmunoResearch (715-155-151) | AB_2340807 |

### Calcium imaging

Segments of mesentery connected to the terminal ileum from Wnt1-Cre;R26R-GCaMP3 mice were used for Ca^2+^ imaging. The mesenteric tissue was isolated, pinned in a silicone elastomer (Sylgard)-bottom dish and then stabilized by an O-ring clamped over an inox ring for imaging (Li et al., 2019). For some experiments, the ileum was opened either adjacent to the mesenteric border or opened along the anti-mesenteric border such that the mesentery remained attached to the gut wall. In the latter case, additional care was taken at points where the vasculature enter the intestinal wall and fine spring scissors were used to sever the vessels traversing the multiple gut layers to minimize damage to the nerve plexus. The mucosa and submucosa were removed by microdissection, and the remaining peeled myenteric plexus-mesentery preparations were stabilized using inox rings. Ring preparations were imaged using a 20x objective on a Zeiss Axiovert 200M microscope equipped with a monochromator (Poly V) and a cooled CCD camera (Imago QE) (TILL Photonics).

All preparations were constantly superfused (approximately 1 ml/min) with Krebs solution (containing in mM: 120.9 NaCl, 5.9 KCl, 1.2 MgCl_2_, 2.5 CaCl_2_, 1.2 NaH_2_PO4, 14.4 NaHCO_3_, and 11.5 glucose, bubbled with carbogen (95% O_2_-5% CO_2_)) at room temperature throughout the experiment.

Electrical stimuli (20 Hz, 2s, 30V) were applied via a stimulating electrode (50 μm tungsten wire). For chemical stimulation, different agonists were locally applied onto neurons and ganglia by pressure ejection (Picospritzer II; 10 psi, 2 sec, General Valve cooperation) using a micropipette (tip diameter of 10-20 µm) containing 5-HT (1 mM), DMPP (100 µM), or ATP (100 µM). For some experiments, we applied high K^+^ (75 mM) Krebs via a perfusion pipette positioned above the myenteric plexus.

For imaging mesenteric neuronal activity in response to ileal distension, the terminal ileum with mesentery attached was collected in a Sylgard-lined dish and the intestinal tube was cannulated at both ends. The mesentery was then stretched and pinned out on Sylgard to restrict movement. Mesenteric neurons were imaged using an M165 FC fluorescent stereomicroscope (Leica) with an X-Cite 200DC Stabilized Fluorescence Light Source (Lumen Dynamics) and fitted with an ORCA-Flash4.0 V3 Digital CMOS camera (Hamamatsu).

For experiments involving an antagonist, ondansetron (Ond; 5-HT_3_ receptor antagonist; 10 µM) or hexamethonium (Hex; nicotinic antagonist; 200 µM) was administrated 10 min before stimulation. Preparations were stimulated first in control Krebs, then in the presence of an antagonist, and followed by a 10 min drug washout period with Krebs.

For experiments examining the connections between the gut and prevertebral sympathetic neurons, we isolated the celiac and superior mesenteric ganglion (CG-SMG) complex (Trevizan-Baú et al., 2024) and the ileum, together with the mesenteric connections between the two. The tissue was then pinned out in a Slygard-lined recording dish. Excess connective tissue and visceral fat was carefully removed with fine spring scissors and forceps, while avoiding cutting the connecting mesenteric nerve fibers. The ileum was then dissected as described above to expose the myenteric plexus. These preparations were imaged using a 10x objective on a Zeiss Examiner microscope (Axio Examiner.Z1; Carl Zeiss) with a monochromator (Poly V) and a cooled CCD camera (Imago QE) (TILL Photonics).

### Drugs

Agonists used included serotonin (5-HT; 1 mM; Kali-chemie pharma Hanover), 1,1-dimethyl-4-phenylpiperazinium (DMPP, nicotinic agonist; 100 µM; Fluka Chemika), and ATP (100 µM; Sigma). Antagonists used include Ondansetron (Ond; 10 µM) and hexamethonium (Hex; 200 µM; Sigma-Aldrich). Drugs were diluted in Krebs to working concentrations at the time of the experiment.

### Analysis

Ca^2+^ imaging analyses were performed using Igor Pro (Wavemetrics, Lake Oswego, Oregon, USA) (Li et al., 2019). Regions of interest were drawn over an area of the cytoplasm of each neuron excluding the nucleus. The amplitude of each [Ca^2+^]i peak was calculated for each response and was measured as the maximum increase in fluorescence from baseline (ΔFi/F0). [Ca^2+^]i transients were considered when the signal increased above baseline by at least 3 times the intrinsic noise. The amplitude of the response evoked in the presence of antagonists was normalized to the corresponding control conditions. The total duration of Ca^2+^ imaging experiments lasted up to 4.5 h following dissection of the tissue out of the mice.

### Data and statistics

The numbers of ganglia and neurons were presented as mean ± SEM. All calcium imaging results are presented as mean percentage ΔF_i_/F_o_ of the control ± SEM. Statistical comparisons between numbers of ganglia and neurons or responses between controls and stimulation conditions were performed using one-way ANOVA with Dunnett’s *post-hoc* test, or a paired t-test where appropriate. P values of < 0.05 were considered to be significant. All statistical analyses were performed using Graphpad Prism.

### Neuronal tracing using DiI or DiD

Some imaged preparations were labelled with DiI (dissolved in 70% ethanol). DiI was locally applied by pressure ejection (Picospritzer II; 10 psi, 2 sec, General Valve cooperation) using a micropipette (tip diameter of ∼10-20 µm) positioned along a mesenteric nerve bundle. To label the mesenteric nerves from the spinal cord side, preparations of peeled ileal myenteric plexus with the mesentery attached on one side were stretched and pinned out in a sylgard dish. Mesenteric nerve trunks were cleaned of adipose and connective tissue before injecting DiD (Invitrogen) (dissolved in 70% ethanol). Tissues were fixed in 4% paraformaldehyde/PBS for 2 h at room temperature. This was followed by PBS washes (3 x 10 min), and tissues were incubated in PBS at 37°C for at least 2 days prior to imaging DiI-labelled preparations, and at least 1 week and up to 28 days post-injection for quantification of DiD-labelled preparations. DiI-labelled preparations were imaged using either a Leica M165 FC fluorescent stereomicroscope (Leica), equipped with an X-Cite 200DC Stabilized Fluorescence Light Source (Lumen Dynamics) and ORCA-Flash4.0 V3 Digital CMOS camera (Hamamatsu), an epifluorescence microscope (BX 41 784 Olympus, Olympus, Aartselaar, BE) with an XM10 (Olympus) camera, and/or a LSM 780 confocal microscope (Zeiss) using a water-immersion 25x objective 1036 (NA 0.8, Zeiss). DiD-labelled preparations were imaged using a Zeiss Axio Imager M2 with an Axiocam 705 (Zeiss).

## Acknowledgements

We thank M. Moons for technical assistance, members of the Laboratory for Enteric Neuroscience (LENS) for advice, and Jean-François Brunet for the Phox2b antibody.

Grant support: METH/014/05, KU LEUVEN. Images were recorded on microscopy equipment in LENS/Cell and tissue Imaging Cluster (CIC) supported by Hercules foundation AKUL/11/37, AKUL/09/50 and FWO G.0929.15.

## References

Boesmans, W., Lasrado, R., Vanden Berghe, P., & Pachnis, V. (2015). Heterogeneity and phenotypic plasticity of glial cells in the mammalian enteric nervous system. Glia, 63(2), 229–241. 10.1002/glia.22746

Bornstein, J. C. (2006). Intrinsic sensory neurons of mouse gut - toward a detailed knowledge of enteric neural circuitry across species. Focus on ‘characterization of myenteric sensory neurons in the mouse small intestine’. Journal of Neurophysiology, 96(3), 973–974. 10.1152/jn.00511.2006

Breunig, E., Michel, K., Zeller, F., Seidl, S., v Weyhern, C. W. H., & Schemann, M. (2007). Histamine excites neurones in the human submucous plexus through activation of H(1), H(2), H(3) and H(4) receptors. J Physiol, 583(Pt 2), 731–742. 10.1113/jphysiol.2007.139352

Coffey, J. C., Byrnes, K. G., Walsh, D. J., & Cunningham, R. M. (2022). Update on the mesentery: structure, function, and role in disease. The Lancet Gastroenterology & Hepatology, 7(1), 96–106. 10.1016/S2468-1253(21)00179-5

Coffey, J. C., & O’Leary, D. P. (2016). The mesentery: structure, function, and role in disease. The Lancet Gastroenterology & Hepatology, 1(3), 238–247. 10.1016/S2468-1253(16)30026-7

Cracco, C., & Filogamo, G. (1993). Mesenteric neurons in the adult rat are responsive to ileal treatment with benzalkonium chloride. International Journal of Developmental Neuroscience, 11(1). 10.1016/0736-5748(93)90034-B

Fung, C., & Vanden Berghe, P. (2020). Functional circuits and signal processing in the enteric nervous system. Cellular and Molecular Life Sciences. 77, 4505–4522 10.1007/s00018-020-03543-6

Fung, C., Venneman, T., Holland, A. M., Martens, T., Alata, M. I., Hao, M. M., Alar, C., Obata, Y., Tack, J., Sifrim, A., Pachnis, V., Boesmans, W., & Vanden Berghe, P. (2025). Nutrients activate distinct patterns of small-intestinal enteric neurons. Nature. 644, 1069–1077 10.1038/s41586-025-09228-z

Furness, J. B. (2003). Intestinofugal neurons and sympathetic reflexes that bypass the central nervous system. Journal of Comparative Neurology 455(3). 10.1002/cne.10415

Furness, J. B., & Costa, M. (1982). Neurons with 5-hydroxytryptamine-like immunoreactivity in the enteric nervous system: their projections in the guinea-pig small intestine. Neuroscience, 7(2), 341–349. 10.1016/0306-4522(82)90271-8

Furness, J. B., Koopmans, H. S., Robbins, H. L., & Lin, H. C. (2000). Identification of intestinofugal neurons projecting to the coeliac and superior mesenteric ganglia in the rat. Autonomic Neuroscience, 83(1), 81–85. 10.1016/S0165-1838(00)00159-4

Lasrado, R., Boesmans, W., Kleinjung, J., Pin, C., Bell, D., Bhaw, L., McCallum, S., Zong, H., Luo, L., Clevers, H., Vanden Berghe, P., & Pachnis, V. (2017). Lineage-dependent spatial and functional organization of the mammalian enteric nervous system. Science, 356(6339), 722–726. 10.1126/science.aam7511

Li, Z., Hao, M. M., Van den Haute, C., Baekelandt, V., Boesmans, W., & Vanden Berghe, P. (2019). Regional complexity in enteric neuron wiring reflects diversity of motility patterns in the mouse large intestine. Elife, 8, e42914. 10.7554/eLife.42914

Lu, H., Feng, Z., Aden, K., Cong, Y., & Liu, Z. (2025). Two sides of the same coin: protective versus pathogenic mesentery in inflammatory bowel diseases – a comprehensive review. International Journal of Surgery, 111(11). 10.1097/JS9.0000000000003075

Mao, Y., Wang, B., & Kunze, W. (2006). Characterization of Myenteric Sensory Neurons in the Mouse Small Intestine. Journal of Neurophysiology, 96(3), 998–1010. 10.1152/jn.00204.2006

Muller, P. A., Matheis, F., Schneeberger, M., Kerner, Z., Jové, V., & Mucida, D. (2020). Microbiota-modulated CART^+^ enteric neurons autonomously regulate blood glucose. Science, 370(6514), 314–321. 10.1126/science.abd6176

Muller, P. A., Schneeberger, M., Matheis, F., Wang, P., Kerner, Z., Ilanges, A., Pellegrino, K., del Mármol, J., Castro, T. B. R., Furuichi, M., Perkins, M., Han, W., Rao, A., Picard, A. J., Cross, J. R., Honda, K., de Araujo, I., & Mucida, D. (2020). Microbiota modulate sympathetic neurons via a gut–brain circuit. Nature, 583(7816). 10.1038/s41586-020-2474-7

Qu, Z. D., Thacker, M., Castelucci, P., Bagyánszki, M., Epstein, M. L., & Furness, J. B. (2008). Immunohistochemical analysis of neuron types in the mouse small intestine. Cell Tissue Res, 334, 147–161. 10.1007/s00441-008-0684-7

Sang, Q., & Young, H. M. (1996). Chemical coding of neurons in the myenteric plexus and external muscle of the small and large intestine of the mouse. Cell Tissue Res, 284(1), 39–53. 10.1007/s004410050565

Schäffler, A., Schölmerich, J., & Büchler, C. (2005). Mechanisms of disease: Adipocytokines and visceral adipose tissue - Emerging role in intestinal and mesenteric diseases. In Nature Clinical Practice Gastroenterology and Hepatology 2(2). 10.1038/ncpgasthep0090

Sheehan, D. (1933). The afferent nerve supply of the mesentery and its significance in the causation of abdominal pain. *Journal of Anatomy*, Pt 2(67), 233–249.

Stebbing, M. J., & Bornstein, J. C. (1994). Electrophysiological analysis of the convergence of peripheral inputs onto neurons of the coeliac ganglion in the guinea pig. J Auton Nerv Syst, 46, 93–105. 10.1016/0165-1838(94)90147-3

Stebbing, M. J., Shafton, A. D., Davey, C. E., Di Natale, M. R., Furness, J. B., & McAllen, R. M. (2024). A ganglionic intestinointestinal reflex activated by acute noxious challenge. American Journal of Physiology-Gastrointestinal and Liver Physiology, 326(4), G360–G373. 10.1152/ajpgi.00145.2023

Szurszewski, J. H., & Linden, D. R. (2012). Physiology of Prevertebral Sympathetic Ganglia. In *Physiology of the Gastrointestinal Tract*, Two Volume Set. 10.1016/B978-0-12-382026-6.00020-8

Tassicker, B. C., Hennig, G. W., Costa, M., & Brookes, S. J. H. (1999). Rapid anterograde and retrograde tracing from mesenteric nerve trunks to the guinea-pig small intestine in vitro. Cell and Tissue Research, 295(3), 437–452. 10.1007/s004410051250

Thomas, E. A., Bertrand, P. P., & Bornstein, J. C. (2000). A computer simulation of recurrent, excitatory networks of sensory neurons of the gut in guinea-pig. Neurosci Lett, 287(2), 137–140. 10.1016/s0304-3940(00)01182-4

Thomas, E. A., & Bornstein, J. C. (2003). Inhibitory cotransmission or after-hyperpolarizing potentials can regulate firing in recurrent networks with excitatory metabotropic transmission. Neuroscience, 120(2), 333–351. 10.1016/s0306-4522(03)00039-3

Timmermans, J.-P., Barbiers, M., Scheuermann, D. W., Stach, W., Adriaensen, D., & De Groodt-Lasseel, M. H. A. (1993). Occurrence, distribution and neurochemical features of small intestinal neurons projecting to the cranial mesenteric ganglion in the pig. Cell & Tissue Research, 272(1), 49–58. 10.1007/BF00323570

Tiveron, M. C., Hirsch, M. R., & Brunet, J. F. (1996). The expression pattern of the transcription factor Phox2 delineates synaptic pathways of the autonomic nervous system. Journal of Neuroscience, 16(23). 10.1523/jneurosci.16-23-07649.1996

Trevizan-Baú, P., Ringuet, M. T., Stebbing, M. J., McAllen, R. M., Furness, J. B., & Mueller, S. N. (2024). Protocol for the isolation of the mouse sympathetic splanchnic-celiac-superior mesenteric ganglion complex. STAR Protocols, 5(2), 103036. 10.1016/j.xpro.2024.103036

Wang, T., Teng, B., Yao, D. R., Gao, W., & Oka, Y. (2025). Organ-specific sympathetic innervation defines visceral functions. Nature, 637(8047), 895–902. 10.1038/s41586-024-08269-0

Yin, Y., Zhu, Z.-X., Li, Z., Chen, Y.-S., & Zhu, W.-M. (2021). Role of mesenteric component in Crohn’s disease: A friend or foe? World Journal of Gastrointestinal Surgery, 13(12). 10.4240/wjgs.v13.i12.1536

Yu, Q., Du, M., Zhang, W., Liu, L., Gao, Z., Chen, W., Gu, Y., Zhu, K., Niu, X., Sun, Q., & Wang, L. (2021). Mesenteric Neural Crest Cells Are the Embryological Basis of Skip Segment Hirschsprung’s Disease. Cellular and Molecular Gastroenterology and Hepatology, 12(1), 1–24. 10.1016/j.jcmgh.2020.12.010

Zhang, T., Perkins, M. H., Chang, H., Han, W., & de Araujo, I. E. (2022). An inter-organ neural circuit for appetite suppression. Cell, 185(14), 2478–2494.e28. 10.1016/j.cell.2022.05.007

